# Model-based characterization of the selectivity of neurons in primary visual cortex

**DOI:** 10.1101/2021.09.13.460153

**Authors:** Felix Bartsch, Bruce G. Cumming, Daniel A. Butts

## Abstract

To understand the complexity of stimulus selectivity in primary visual cortex (V1), models constructed to match observed responses to complex time-varying stimuli, instead of to explain responses to simple parametric stimuli, are increasingly used. While such models often can more accurately reflect the computations performed by V1 neurons in more natural visual environments, they do not by themselves provide insight into established measures of V1 neural selectivity such as receptive field size, spatial frequency tuning and phase invariance. Here, we suggest a series of analyses that can be directly applied to encoding models to link complex encoding models to more interpretable aspects of stimulus selectivity, applied to nonlinear models of V1 neurons recorded in awake macaque in response to random bar stimuli. In linking model properties to more classical measurements, we demonstrate several novel aspects of V1 selectivity not available to simpler experimental measurements. For example, we find that individual spatiotemporal elements of the V1 models often have a smaller spatial scale than the overall neuron sensitivity, and that this results in non-trivial tuning to spatial frequencies. Additionally, our proposed measures of nonlinear integration suggest that more classical classifications of V1 neurons into simple versus complex cells are spatial-frequency dependent. In total, rather than obfuscate classical characterizations of V1 neurons, model-based characterizations offer a means to more fully understand their selectivity, and provide a means to link their classical tuning properties to their roles in more complex, natural, visual processing.

**Significance statement:** Visual neurons are increasingly being studied with more complex, natural visual stimuli, with increasingly complex models necessary to characterize their response properties. Here, we describe a battery of analyses that relate these more complex models to classical characterizations. Using such model-based characterizations of V1 neurons furthermore yields several new insights into V1 processing not possible to capture in more classical means to measure their visual selectivity.

## Introduction

The responses of neurons in primary visual cortex (V1) set the foundation for the cortical processing of vision. Thus, detailed characterization of their stimulus selectivity is of great intrinsic utility. Since the initial discovery of their basic selectivity to oriented bars (Hubel and Wiesel, 1962b), most V1 neurophysiology has used precise manipulations of simple parametric stimuli such as flashed or drifting gratings (Movshon et al., 1978a; Maunsell and Newsome, 1987; Alonso and Martinez, 1998) to make measurements of dominant aspects of V1 visual selectivity, such as receptive field (RF) size and location, orientation tuning, and spatial frequency and temporal frequency tuning (Hubel and Wiesel, 1962a; DeAngelis et al., 1994; Sceniak et al., 1999; Chen et al., 2009). Because how V1 processes these simple parametric stimuli has limited power in predicting their responses to more complex time-varying stimuli (David and Gallant, 2005; Sharpee, 2013; Butts, 2019), a large number of increasingly sophisticated models has been used to understand their responses to more complex stimuli (Butts, 2019), using techniques ranging from spike-triggered covariance (Touryan et al., 2002; Rust et al., 2005) up through modern machine-learning methods (e.g. (Cadena et al., 2019)). While the necessity of more sophisticated models to capture neural computation in complex stimulus contexts suggests that these previous measures of V1 selectivity offer an incomplete picture of its processing of natural vision, it is not clear whether more sophisticated approaches are simply measuring different aspects of processing, and exactly what is gained by them relative to more canonical measurements.

Indeed, the success of more sophisticated models in explaining V1 neural responses is often at the expense of the interpretability that defined more canonical approaches to characterizing V1 neurons (Butts, 2019). Likewise, few studies analyzing models to gain insight into neural properties have attempted to directly relate modern nonlinear modeling approaches to classical measures of cell (although see (Almasi et al., 2020)). Here, we focus on the most interpretable of these modern models: “subunit” models such as the LNLN cascade (Park and Pillow, 2011; McFarland et al., 2013; Vintch et al., 2015), which consist of a linear combination of nonlinear spatiotemporal stimulus filters (or “subunits”) that are each selective to a particular stimulus feature that contributes to the neuron’s response (Vintch et al., 2012, 2015; McFarland et al., 2013; Park et al., 2013). The array of spatiotemporal filters generated by these models are determined by optimization of model parameters using data recorded during any sufficiently general spatiotemporally varying visual stimulus paradigm, and captures the set of stimulus features that affect the neuron’s response. We present several measures that can then be derived from the resulting models, which can be directly related to canonical aspects of neural tuning, while capturing other interpretable aspects of its selectivity. Notably, while we apply these measures to the models measured in this particular context, such measures can be generalized to any “image-computable” model of V1 neurons (Butts, 2019). Because such more sophisticated characterizations of V1 processing implicitly capture V1 neural selectivity to multiple features as well as interactions between selectivity to related features, the resulting descriptions of V1 computation go beyond what can be achieved with simple parametric stimuli, and set a new bar for measurements of V1 neural selectivity.

## Materials and Methods

### Neurophysiology experiments

We used previously collected recordings, as described in detail in (McFarland et al., 2014). Briefly, multi-electrode recordings were made from primary visual cortex (V1) of two awake head-restrained male rhesus macaques (*Macaca mulatta*; 13- to 14-years old) while the animals performed a simple fixation task, where they were required to maintain gaze within a small window around a fixation target to obtain a liquid reward after each completed 4-s trial. Eye position was monitored continuously using scleral search coils in each eye, sampled at 600 Hz, and a model-based eye-position correction algorithm (see below) was applied to accurately recover eye position at sufficiently high spatial resolution (McFarland et al., 2014, 2016). Trials during which the monkey broke fixation were discarded from analysis. Additionally, the first 200 ms and last 50 ms of each trial were excluded from analysis to minimize the impact of onset transients.

### Recordings

In one animal, we implanted a 96-electrode planar Utah array (Blackrock Microsystems; 400 mm spacing), in the other, a linear electrode array (V-probe, Plexon; 24 contacts, 50 μm spacing) was passed through the dura at the start of each day. Spike waveforms were detected online using Spike2 software (Cambridge Electronic Design) and recorded at 32 kHz (Utah array) or 40KHz (V-Probe) sampling rate. Single-units (SUs) were identified through off-line spike-sorting with custom software. Spike clusters were modelled using Gaussian mixture distributions that were based on several different spike features, including principal components, voltage samples and template scores. The features providing the best cluster separation were used to cluster single units. Cluster quality was quantified using a variety of measures including ‘L-ratio’, ‘isolation distance’, as well as a variant of ‘d-prime’. Only spike clusters that were well isolated using these measures—confirmed by visual inspection— and had average firing rates during stimulus presentation of at least 5 spikes per second were used for analysis. Voltage signals were also separately low-passed at 100 Hz and used to compute the inverse current-source density (iCSD) profile of each recording, in order to categorize each isolated single unit by layer (Chen et al., n.d.; Pettersen et al., 2006). All protocols were approved by the Institutional Animal Care and Use Committee and complied with Public Health Service policy on the humane care and use of laboratory animals.

### Stimuli

We presented a ternary bar noise stimulus, consisting of random patterns of black, white, and grey bars (matching the mean luminance of the screen), centered on the neurons’ RFs. These stimuli were displayed on cathode ray tube (CRT) monitors at a 100 Hz refresh rate, and subtended 24.3°×19.3° of visual angle. Individual bars had a width of 0.057° for the Utah array recordings and ranged from 0.038° to 0.1° for the linear array recordings, depending on the RF sizes of the recorded neurons. Bar patterns uncorrelated in space and time. The probability of a given bar being non-grey (i.e., black or white) was set to 88% grey (relatively sparse), although in several experiments we used a denser distribution (33% grey), which yielded similar results. For the linear array recordings, the orientation of the bars was chosen to be within 15 degrees of the preferred orientation of the majority of units recorded in a given session. In Utah array recordings, where the neurons typically had a full range of preferred orientations due to being located across multiple orientation columns, we performed the experiments with two different bar orientations (vertical and horizontal). In this case we modeled the orientation that best drove neural activity, and the properties of cells recorded using this approach were comparable to those recorded during presentation of more optimally oriented bar stimuli.

### Model-based eye tracking

Due to fixational eye movements, the stimulus projected on the retina is shifted over time, which can result in mis-estimation of model parameters (McFarland 2014; 2016). As a result, we estimated the precise position of the eye as a function of time using model-based eye tracking (McFarland et al., 2014). Briefly, the algorithm uses the observed firing rates during each fixation period to estimate the probability of each potential eye position, and — with enough neurons simultaneously recorded — these probabilities can be combined to accurately infer the most likely eye position, which can be accurate to arc-min resolution. Using this estimated position, we then adjusted the presented stimulus by the eye shifts, and used the resulting adjusted stimulus to fit the models, as described in detail in (McFarland et al., 2014).

### Model structure

We used the Nonlinear Input Model (NIM) to model V1 neurons (McFarland et al., 2013). The NIM (Fig. 1A) operates on the stimulus **s**(*t*) with a number of linear-nonlinear (LN) subunits, each consisting of a stimulus filter **k** (Fig. 1B) and a rectifying nonlinearity *f*(.). Subunit outputs are then weighted by either +1 or -1 (depending on whether each is excitatory or inhibitory, respectively), and summed together and and passed through a final spiking nonlinearity *F*[.] to result in a predicted firing rate *r*(*t*): given stimulus **s**(*t*) is given as:

**Figure 1:**
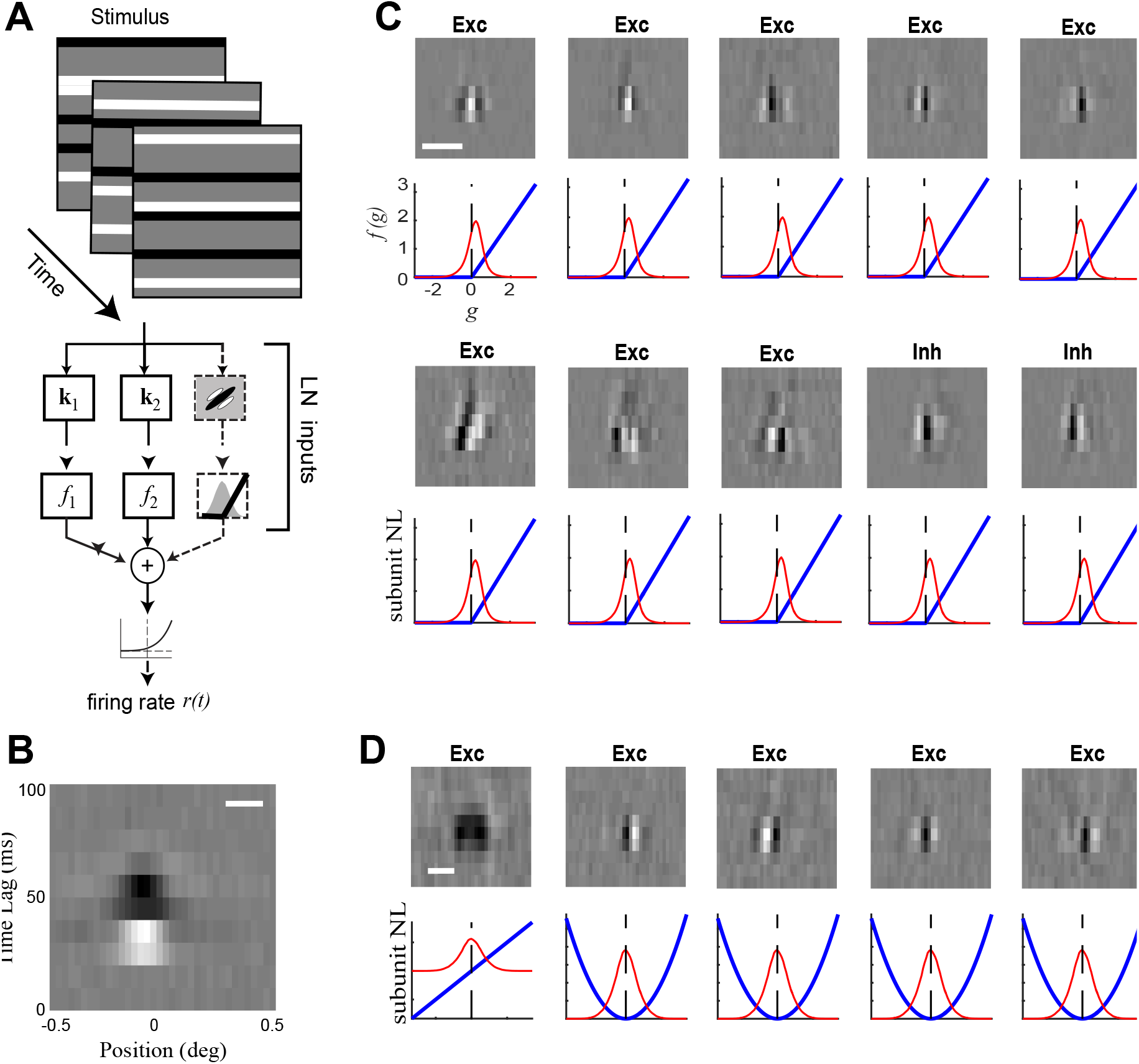
Extracting RF components using stimulus models. ***A***, Schematic of the nonlinear LNLN cascade stimulus processing models. Stimulus frames (top) are passed through spatiotemporal filters **k**_*i*_, the outputs of which are then passed through a rectifying nonlinearity. The outputs of these “LN subunits” is then summed to give the ‘generating signal’ *g*(*t*), which is transformed into a firing rate by the spiking nonlinearity (bottom). In the case of the PCNIM, filters **k**_*i*_ were not fit using the full stimulus space, but rather its low-dimensional projection into the space outlined by the principal components. After fitting, they were retranslated into full space for interpretation. ***B***, Example spatiotemporal filter. ***C***, Example of filters fit by a GQM architecture with 4 excitatory quadratic filters. The nonlinearity associated with each filter is shown below (blue). ***D***, Example of filters fit using the PCNIM with 8 excitatory and 2 suppressive filters. Nonlinearities are again displayed below each filter in blue. The scale bars in panels B-D indicate 0.1 degrees of visual angle.

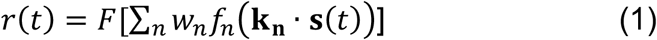

where *w*_*n*_ is +1 or -1 as described above.

We initially fit NIMs to each neuron using standard maximum *a posteriori* optimization in the context of sparseness (L1) and smoothness (Laplacian) regularization. Both the number of subunits and the regularization parameters were optimized through nested cross-validation, as described previously (McFarland et al., 2013; Butts, 2019). We then leveraged the resulting collection of models to yield additional model improvements, using dimensionality reduction on the spatiotemporal filters pooled across all models. Specifically, this database (over all neurons in the dataset fit using standard NIMs) was consisted of 2061 filters. We computed the principal components (PCs) of this filterbank, finding that 80% of its variance could be explained by the first 50 PCs, with diminishing returns from including more (see Appendix Figure A1A). We then applied these top 50 filters to the spatiotemporal stimulus **s**, resulting in a 50-dimensional “dimensionality-reduced” stimulus in place of the 360-dimensional “full” stimulus that was composed of 36 spatial dimensions and 10 lags.

The resulting dimensionality-reduced stimulus was used in eq. 1 in place of the full stimulus, and NIMs were refit using the filtered, dimensionality-reduced stimulus. The fit model subunits thus had a set of coefficients corresponding to the weights on the PCs, and the full spatiotemporal filters could be recovered by matrix multiplication. This allowed for the potential inclusion of many more subunits without overfitting (see below). Furthermore, because the PCs did not reflect noise in the filter estimation that was not shared among many filters, it was not necessary to apply any regularization on subsequent stages of fitting. Representing model filters using PCA yielded better model performance (Appendix Fig. A1C) than the standard NIM approach, including finding more subunits that improve model performance that would have been suppressed by regularization penalties for the standard NIM.

In addition to using the standard structure of the NIM with rectified subunits, we also compared LNLN cascades with linear and quadratic subunit nonlinearities *f*_*n*_(.) (eq. 1) (McFarland et al., 2013; Park et al., 2013; Park & Pillow, 2011) (Fig. 1C). This “quadratic” model is also commonly used to model V1 neurons because it embodies the “energy model”(Adelson and Bergen, 1985a; Mante and Carandini, 2005), although less flexible in approximating more general nonlinearities (McFarland et al., 2013; Butts, 2019).

### Model parameter optimization

The parameters of the filters are fit and optimized through a maximum-likelihood approach as described in detail (McFarland et al., 2013). This approach maximizes the log-likelihood of the model, which is given by the equation

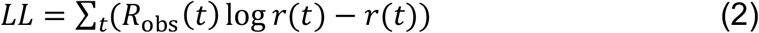

where *r(t)* is the predicted firing rate and *R*_obs_(*t*) is the observed spike count. The goodness of fit of the models is assessed using 5-fold cross-validation: with model parameters fit using 80% of the data, and performance assessed using the *LL* of the models on the remaining 20% of the data.

### Model hyper-parameter optimization

In addition to the parameters corresponding to the spatiotemporal filters, each model had several “hyper-parameters” corresponding to other aspects of the model architecture, including the number of filters used and the ratio of excitatory to inhibitory filters (E-I ratio). The number of filters and E-I ratio were determined through nested cross-validation across a range of possible combinations, refitting model parameters for each new combination and choosing the combination yielding the best cross-validated performance. Note that because subunits were fit in a subspace, they inherited the regularization of the models that were used to generate the subspace (see above).

### Estimating spatial frequency tuning via forward correlation

To compare model-based measures of spatial frequency (SF) tuning to a model-free measure, we used forward correlation. We first extracted spatial frequency spectra of each stimulus frame via fast-Fourier transform (FFT) and ranked frames by power in each resulting frequency bin. For each frequency, we then extract two peri-stimulus time histograms (PSTHs) corresponding to frames with power at that frequency in the top 30^th^ percentileand in the bottom 30^th^ percentile. The difference between these two PSTHs provides a model-free estimate of the neuron’s SF tuning directly comparable to those obtained using classical measures of SF tuning. We defined the cell’s preferred frequency as the peak of this curve.

## RESULTS

### Estimation of a neuron’s spatiotemporal feature space

In order to characterize the stimulus processing of V1 neurons, we fit the Nonlinear Input Model (NIM) (McFarland et al., 2013) to a database of 122 V1 neuron responses recorded in the context of noise stimuli consisting of ternary bars aligned to preferred orientation (Fig. 1A) (McFarland et al., 2013, 2014) (see Methods). The resulting subunit model of a given V1 neuron provides a list of spatiotemporal stimulus filters that modulate the neuron’s response, as well as a positive or negative weight corresponding to whether the feature was excitatory or suppressive (Fig. 1B). As described above, these model components (Fig. 1C) contain rich information about the selectivity of the neuron being modelled, and here we are interested in analyzing the properties of these filters to extract such information.

We first analyzed the spatial profile of the filters: computing the amount of power (i.e., variance across time lag) at each spatial position, which defines the spatial profile for each spatiotemporal filter (Fig. 2A, B). The overall spatial profile of the neuron is then defined as the average of its filter profiles, weighted by the relative contribution of each filter’s output to the overall predicted firing rate (Fig. 2B, bottom panel). To associate a single “width” to a given spatial profile, we defined the width of a given profile using the spatial extent between 30% and 70% of the cumulative spatial power are under the profile curve.

**Figure 2:**
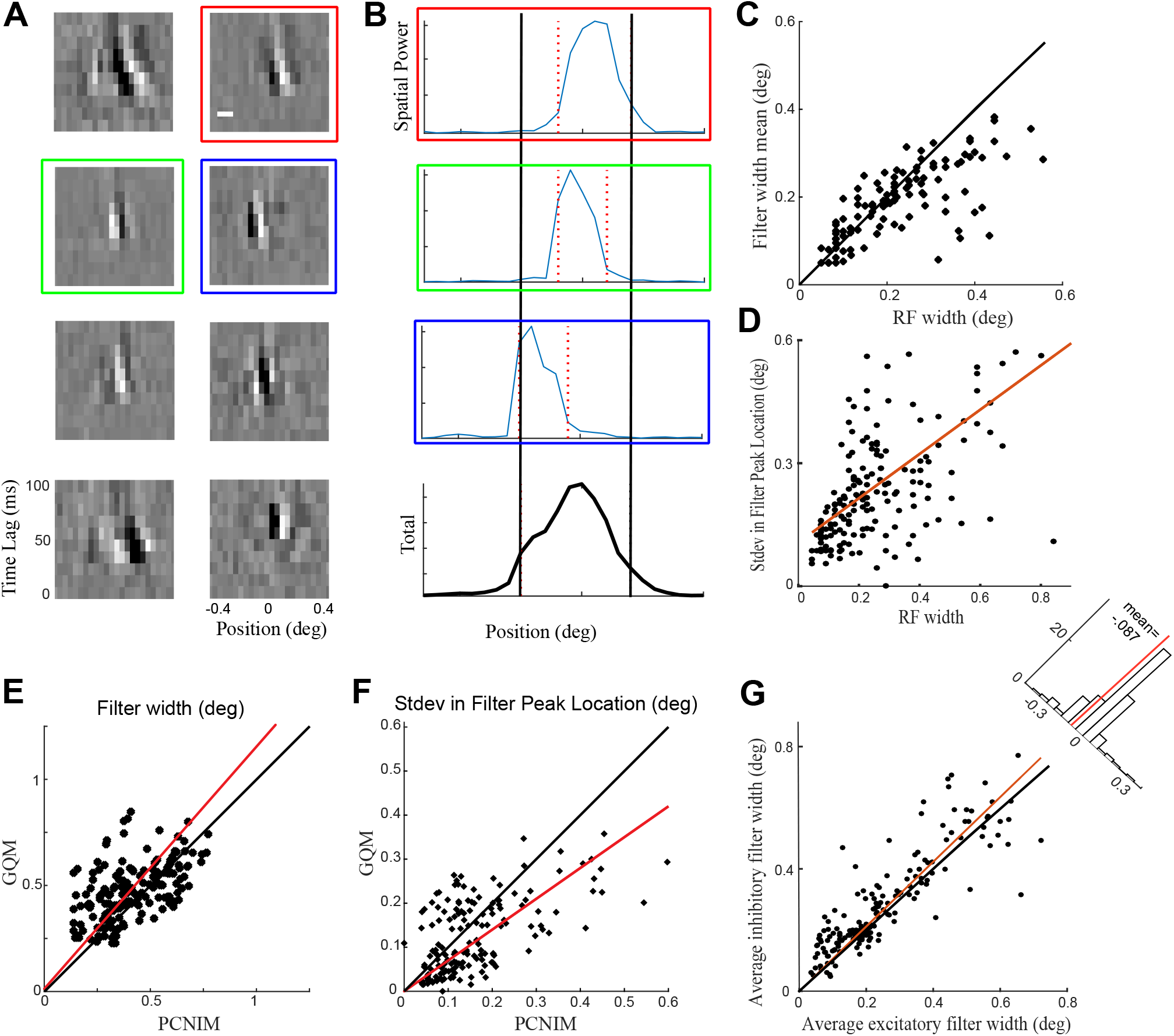
PCNIM reveals high-resolution RF components. ***A***, Example model filters. The horizontal axis denotes spatial position, while the vertical axis shows temporal differences. The eight most highly weighted filters contributing to the model’s predictions are shown in order. ***B***. Model spatial power profiles. Colored boxes indicate which filter’s total spatial power profile (power at each time lag weighted by the filter’s temporal profile) is shown. Vertical lines indicate the edges of each filter, used to determine filter width. The bottom panel shows the sum of filter spatial power, where each filter’s spatial power profile is weighted by its contribution to the model’s overall firing rate prediction. ***C***, Population results comparing RF width and mean filter width. ***D***, Population results comparing RF width and standard deviation of filter locations in excitatory filters. As RF width increases, there is more spatial scatter in the locations of individual filters. ***E***, Filter width compared for GQM and PCNIM fits. The GQM on average fits wider filters. ***F***, Comparing filter location STDV between GQM and PCNIM fits. The GQM generally finds less variability in filter locations, indicating that our highly localized results would be more difficult to identify with a GQM approach. ***G***, Average width of each model’s excitatory and inhibitory filters. Inhibitory filters are overall wider than excitatory filters.

This analysis reveals that, in a large fraction of neurons, there was a significant difference between extent of each neuron’s summed spatial profile and those of its individual subunits. Across the population, a significant fraction of cell RFs are composed of filters with a mean width less than 75% of the total RF width (Fig. 2C). This discrepancy can be explained by a scatter of the locations of a given neuron’s filters (Fig. 2B), such that the full profile of the neuron combines the smaller spatial extent of its component subunits and the scatter of these elements. This finding is not readily accessible using direct experimental measurements of RF spatial potential and scale, which produce a single scale and location or preferred phase (Carandini et al., 1997; Kagan et al., 2002; Mante and Carandini, 2005; Touryan et al., 2005). Notably, spatiotemporal processing at the locations covered by different subunits was often not identical.

Because individual subunits in the NIM are rectified, subunits are implicitly either excitatory (only generating positive or zero output) or suppressive (with a negative weight so its output can only be less than or equal to zero) (McFarland et al., 2013). Such computational distinctions are difficult to assess using classical characterization methods, and reveal systematic differences in the selectivity of putative excitatory and inhibitory elements. For example, suppressive subunits typically had an increased width compared with excitatory subunits (Fig. 2D). This finding is consistent with intracellular recordings that found that the tuning of onto V1 neurons typically have broader spatial profiles (Haider et al., 2012).

To see if our observations were dependent on the specific form of the model, we applied the same analysis to quadratic models fit to each neuron (see Fig. 1C for an example model) (Park and Pillow, 2011; McFarland et al., 2013, 2014). Measurements we applied to the quadratic models yielded the same overall trends for filter width (Fig. 2E) and scatter in subunit location (Fig. 2F). However, the quadratic model filters were on average wider, and the detected scatter was of smaller magnitude. This could be explained by the quadratic model’s use of quadrature pair filters to approximate spatial invariance of the neuron’s selectivity (Lochmann et al., 2013; Almasi et al., 2020). Because the quadratic models also underperformed the NIM at predicting neural responses (Fig. A1C), we take this as indication that one aspect of the increased flexibility with the NIM as a general function approximator (Kriegeskorte, 2015; Butts, 2019) is its ability to identify more spatially localized feature selectivity, and that this more localized feature selectivity allows us to more clearly see trends present in the quadratic models.

### Measuring spatiotemporal selectivity

A dominant conceptual model of V1 neuron selectivity is that is extracts local information about the spatiotemporal content of the stimulus (Adelson & Bergen, 1985; Mante & Carandini, 2005), and a main reason classical tests of V1 tuning involve using measurements of responses to gratings. Here, we can instead extract information about such tuning by Fourier analysis of the model subunits.

Applying two-dimensional Fourier transforms to each subunit (Ringach and Shapley, 2004) yields a two-dimensional power frequency spectrum (Fig. 3A). By summing along either axis of this spectrum, we generate a tuning curve demonstrating the subunit’s tuning preferences to either spatial or temporal frequency, where peak of the subunit spectrum determines its preferred frequency. We can then determine the model’s overall frequency tuning by computing the average tuning spectrum across subunits (again weighted by each subunit’s contribution to the model output) and measuring the peak of this model tuning spectrum (Fig. 3B). The resulting tuning measures can then be compared to classical measures from these models, and also allows for more in-depth analysis of the tuning properties of the cell’s inputs (see below), such as characterizing any variability in the tuning of individual subunits compared to the cell’s overall tuning curve.

**Figure 3:**
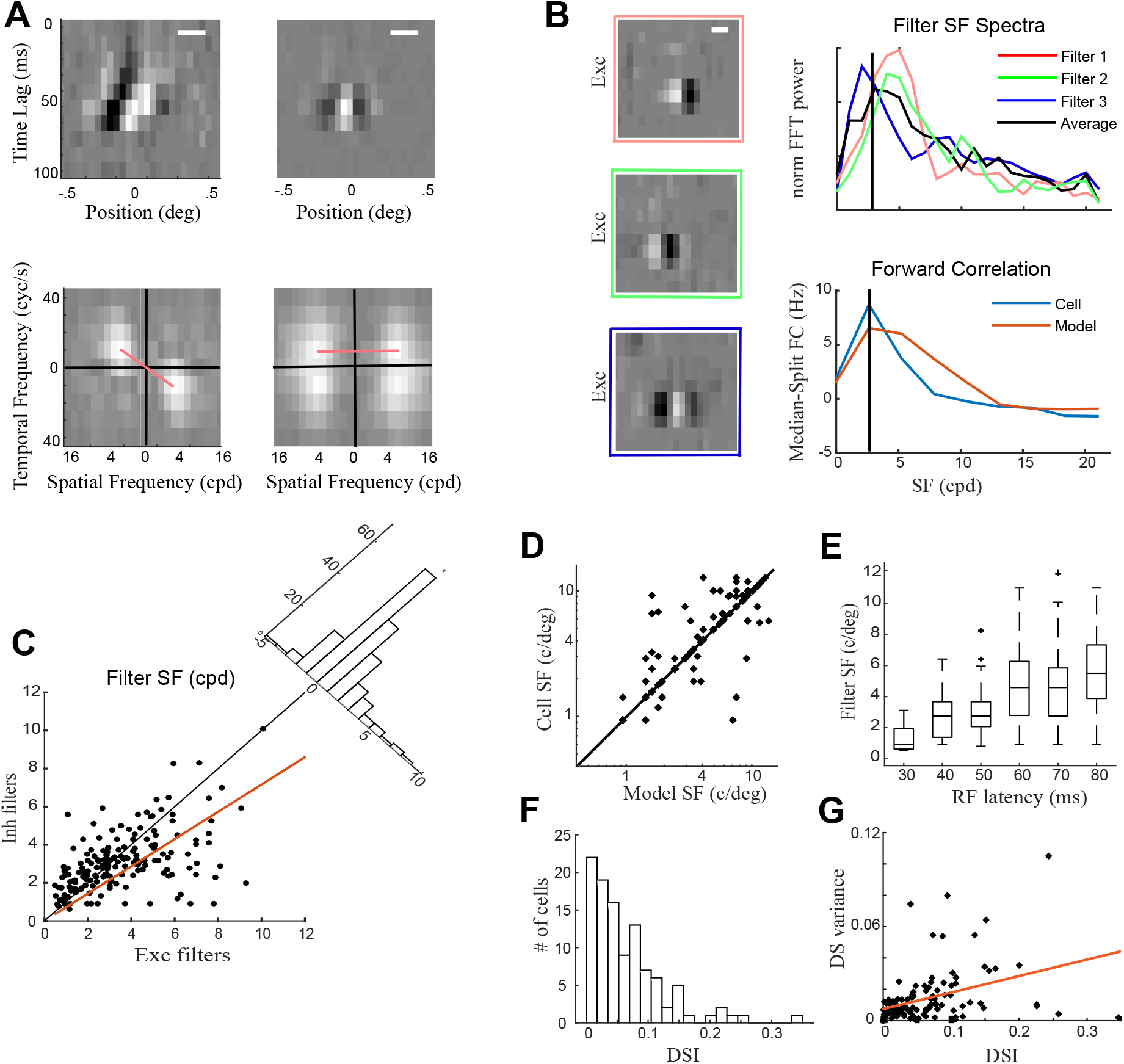
Extracting spatial, temporal, and spatiotemporal feature selectivity from models. ***A***, *Top*: Example filters demonstrating direction selectivity (left) and a non-direction selective filter (right). Scale bars indicate .2 degrees of visual angle. *Bottom*: 2DFFT spectra of the above filters. Peaks in the spatial and temporal dimensions are extracted as a measure of frequency tuning preference of the filter. A slope can be fit to the peak locations of the 2DFFT spectrum (magenta line); the reciprocal of this slope indicates the DS velocity. Additionally, the difference in total power in the tope left and top right quadrants of the 2DFFT spectrum can be used to quantify the DSI. This measure yields both the strength and direction the cell is selective to. ***B***, *Left Column*: Example filters. *Right Column, Top*: SF spectra extracted from the 2DFFT of each filter shown to the left. Each filter tuning curve corresponds to the filter in the left column outlined with the same color. The weighted mean across all filters (weighted by the filter contribution to the model output) is shown in black. *Right Column, Bottom*: Frequency spectrum of the same cell estimated using forward correlation. Black vertical lines indicate the peak frequency. ***C***, Differences in spatial frequency tuning of excitatory and inhibitory filters across our population. Excitatory filters are on average tuned to higher spatial frequencies. ***D***, Preferred spatial frequency as estimated by the model-free forward correlation using observed firing rate and the model prediction. The two measures highly agree, further validating our model approach. ***E***, Filter spatial frequency preference binned by the filter’s onset latency. There is a positive correlation between filter onset delay and spatial frequency tuning. ***F***, histogram of DSI values across our sample. ***G***, variability in DSIs of each filter of a given model VS the model’s overall DSI.

To validate the model-based measures with experimental data – in the absence of direct measurements of the classical tuning curves using gratings -- we used a model-free estimation of spatial frequency tuning using forward-correlation of the spike train of each cell (Spratling, 2012; Tanabe and Cumming, 2014), where we extracted the spatial frequency components of each stimulus frame and identify the frequency components most correlated with the observed neural response (see methods). Applying this procedure to both the observed and the model-predicted firing rates, we found that the two agree well (Fig. 3D), indicating that the models are successfully capturing the cell’s SF tuning.

Model-based characterization also can address how SF tuning relates to other properties in our models. For example, inhibitory subunits generally have selectivity to lower SFs, in line with previous results indicating that inhibitory inputs into V1 tend act on broader spatial scales (Ringach et al., 2003; Haider et al., 2012; Taylor et al., 2021). Additionally, we found a positive relationship between subunit SF selectivity and latency (Fig. 3E), consistent with (Mazer et al., 2002), indicating that low-frequency features have lower onset latencies and thus may be processed faster.

Similarly, we inferred V1 neuron direction selectivity (DS) from our models by extracting the direction selectivity index (DSI) (Priebe and Ferster, 2005) from each filter’s 2DFFT (see Fig. 3A, red lines in bottom panels). We found that subunits of the same model exhibit a variety of spatial frequency sensitivities and DS strengths (Fig. 3G), suggesting that some direction-selective cells will be modulated by stationary inputs. We additionally tested for relationships between DS strength and other tuning properties including SF selectivity and RF size but did not find significant relationships.

### Model-based characterization of SF tuning

Having established model-based measures of spatiotemporal selectivity, we can now compare to more classical ways of estimating SF tuning using drifting grating stimuli (Victor et al., 1994; Ninomiya et al., 2012) to learn how selectivity to SF might be generated. One advantage of model-based approaches is that the models-are “image-computable”: meaning that we can simulate the responses of the model to arbitrary stimuli. While our dataset did not include recordings of responses to drifting gratings, we used the models of our sample cells to simulate their firing rates in response to static and drifting gratings and measured the F0 (mean) and F1 (modulation depth) components of the response to each grating frequency (Fig. 4B, solid lines show the responses of the model displayed in Fig. 4A, left) and compared those to our purely model-based estimates of SF preference. This led to a startling revelation: for some models, the predicted preferred SF tuning in the context of gratings that was to a much lower frequency than that predicted by the model. Closer inspection of these cells’ SF tuning curves revealed two peaks both in their F0 and F1 spectra: with the low-frequency peak in the F1 spectrum indicating the highest response. Normally, the low-SF F1 peak being the largest response would be interpreted as indicating a simple cell tuned to a low SF. However, the F1 outputs of individual filters showed that all were preferentially tuned to higher frequencies (Fig. 4C), indicative of a more complex-like cell that could detect high-frequency inputs. Plotting the outputs in response to different gratings over time revealed that for high-frequency gratings: individual filters were being driven most strongly, but that the output of each subunit did not sum temporally due to their different spatial locations. In contrast, to low-frequency gratings, the relatively small responses of each subunit did sum (Fig. 4D). Therefore, the apparent selectivity to a low-SF grating was in fact generated as a result of a low-SF “envelope” driving multiple high-SF features suboptimally, but simultaneously. Thus, this type of cell appears to be simple-like for low SFs, but complex for high SFs.

**Figure 4:**
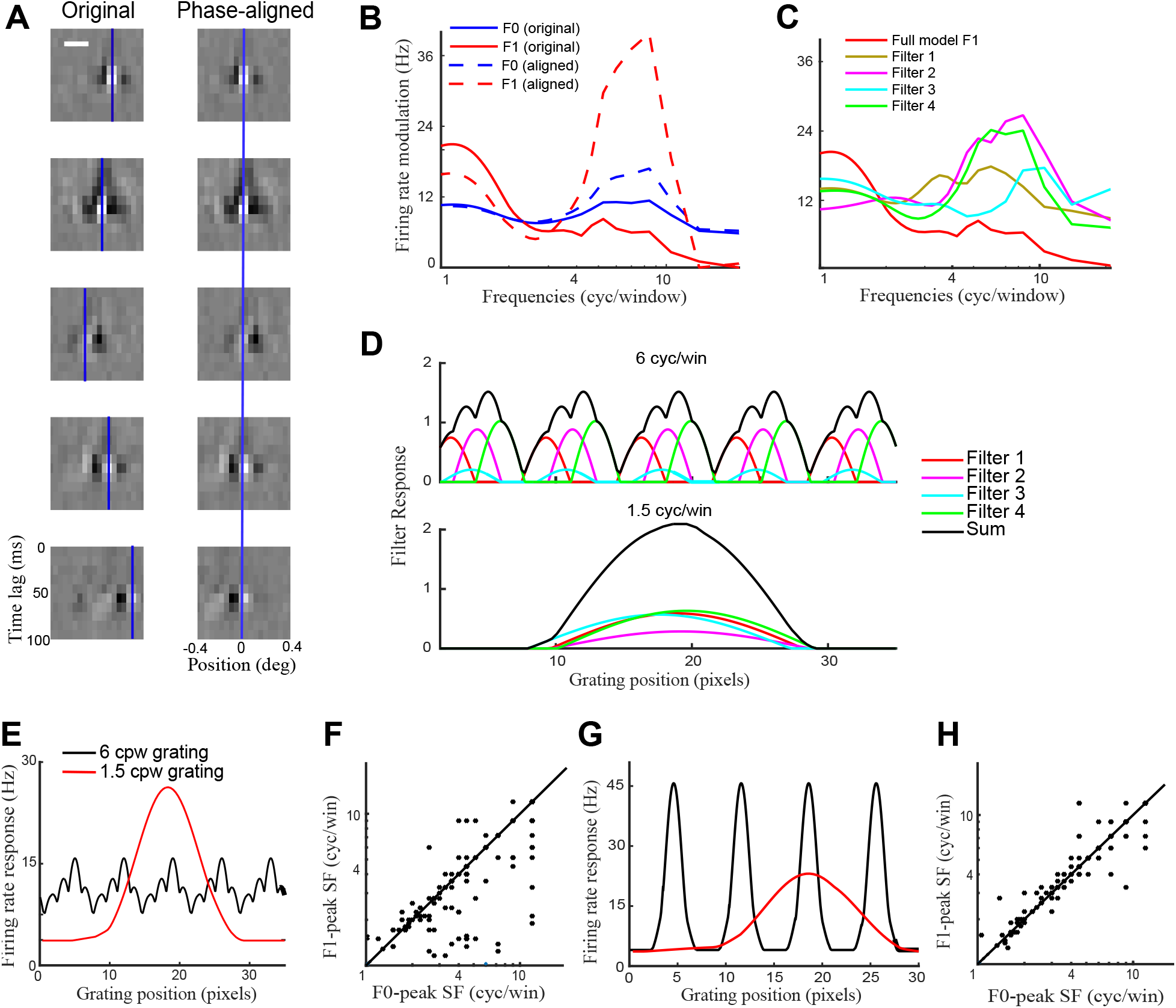
Responses to gratings by individual filters and the whole model. ***A***, Example model of a cell displaying envelope tuning. Full spatiotemporal filters are displayed on the left, while the same filters following phase alignment are displayed on the right. White vertical lines indicate the positive phase peak of the Gabor function used to fit the filter. B, F0 (blue) and F1 (red) responses to simulated gratings for the original and aligned model’s simulated responses to grating stimuli. ***C***, F1-based SF tuning curves of individual filters and the whole cell estimated using simulated responses of the model to standard grating stimuli. ***D***, Responses over time to a high-frequency (HF) 6 cpd grating (top) and a low-frequency (LF) 1.5 cpd grating (bottom) by individual filters (colored lines) and their sum (black line). The HF grating drives individual filters well, but the cell’s overall phase invariance means that the individual responses do not sum, whereas the LF grating at the envelope frequency drives all filters simultaneously. ***E***, Original model output in response to LF and HF gratings over time. The sum of weaker modulation by the LF grating generates a larger response than the individual responses of each filter to the HF grating. ***F***, SF measurements based on F1 and F0. Measuring the grating responses using F0 and F1 reveals that F0 predicts tuning to higher frequencies. ***G***, Original model output in response to LF and HF gratings over time for the phase-aligned model. HF responses sum temporally, the HF response is now again much larger than the LF envelope response. ***H***, SF measurements based on F1 and F0 after phase-alignment. Across the population, this also aligns the preferred SF as estimated using F0 and F1 measures.

Indeed, this apparent discrepancy between subunit tuning and the tuning to gratings can be explained by the previously observed spatial scatter between filters. To demonstrate this, we simulated two models for each neuron: the original model of the data, along with a second model constructed with identical subunits, except that the filters of each subunit were spatially aligned to have the same position, eliminating their spatial scatter (Fig. 4A, *right*). As expected, these peak-aligned models display greatly enhanced tuning to the carrier frequency (Fig. 4B, dotted lines) relative to the original models, and furthermore eliminated the discrepancy between model- and grating-estimated SF tuning (Fig. 4F, H). In the context of classical F0/F1 measures, these results indicate that some cells may show large modulation in their F1 component at a lower frequency, while still having significant tuning to a higher SF that is obscured by the choice of gratings used, similar to some previously observed properties of complex cell responses (Movshon et al., 1978b). Thus, we interpret tuning at the envelope frequency as not being caused by the presence of an explicit envelope filter, but rather as arising from the interactions between individual, rectified, high-resolution filters.

### Quantifying the degree of nonlinearity across cortical layer

Model-based measures can also be useful to identify the degree that a given V1 neuron is nonlinear in its spatiotemporal processing. Traditionally, the degree of nonlinearity of a given V1 neuron has been cast in terms of the simple/complex cell spectrum (Hubel and Wiesel, 1962b; Skottun et al., 1991; Mechler and Ringach, 2002), such characterizations can be undermined by the fact that the degree of nonlinearity a given cell demonstrates will depend on stimulus context (Ringach and Shapley, 2004; Yeh et al., 2009; Butts, 2019; Almasi et al., 2020). Model-based characterizations of nonlinearities thus provide an opportunity to produce more general measurements of the degree of nonlinearity in stimulus processing, which we can then relate back to the simple/complex transformation.

We propose two measures: a holistic measurement of the degree to which the neuron is nonlinear, and a measure that analyzes the nonlinearity as a function of spatial position. To identify the overall degree of nonlinearity throughout the RF, we utilize our previously-published phase-reversal modulation (PRM) measure (McFarland et al., 2014), which more closely corresponds to the classical grating-based F1/F0 measure of phase-invariance. We then expand upon the PRM to include the degree of phase-reversal modulation across the spatial dimension of the stimulus (Fig. 5B). This spatial PRM, or sPRM, is defined as the average difference between the simulated responses to a stimulus and a stimulus where information at one spatial position *i* is inverted, normalized by the overall average rate, and is given by:

**Figure 5:**
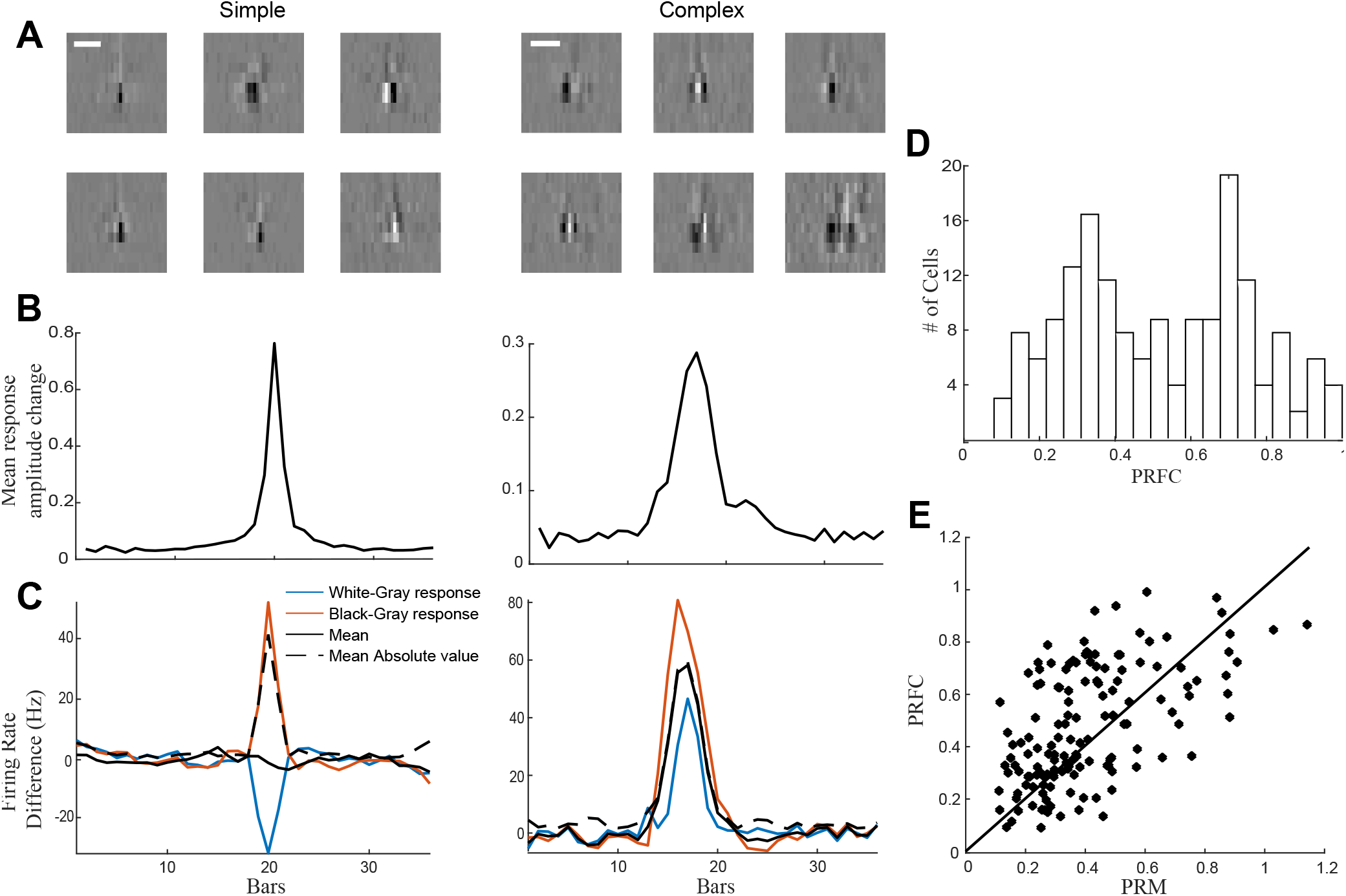
Measures of cell complexity and nonlinearity. ***A***, Example filters of a simple cell (left) and a complex cell (right). ***B***, Spatial PRM profiles for the above cells. These spatial profiles closely match those established by the model filters in B. ***C***, The spatial profiles of the forward-correlation based measure of phase insensitivity (PRFC) corresponding to the above example cells. Triggering on each instance of a bar at one spatial position being non-gray yields a measure of firing rate modulation across space. For a simple cell (left), responses to bars of the opposite sign are also opposite, leading them to average out relative to the average of their absolute values. In a complex cell (right), any non-gray bar increases the firing rate, leaving the average modulation identical to the average of its absolute values. ***D***, Histogram of PRFC values, showing the broadly bimodal distribution normally associated with the simple/complex spectrum. ***E***, Agreement between model-based and model-free measures of cell complexity.

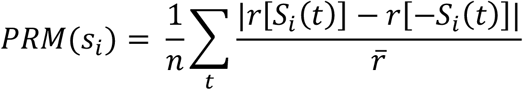

where *r*[**S**_**i**_(*t*)] is the predicted response to the stimulus at time *t, r*[−**S**_**i**_(*t*)] is the predicted response to the polarity-reversed stimulus and 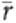 is the average response rate.

To validate these measurements in a model-free way, we use forward-correlation to estimate each neuron’s phase invariance. This phase-reversal forward correlation (PRFC), simply a correlation between each presentation of each bar color and the subsequently observed response, is given by:

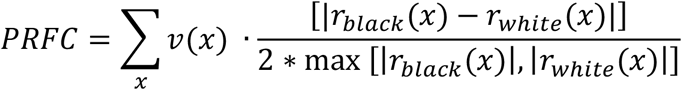

where *x* is each spatial position, *v*(*x*) is the spatial profile of the cell (as established earlier) used to normalize the modulation at each spatial position, *r*_*black*_(*x*) and *r*_*white*_(*x*) are the mean firing rate response when a bar at a given spatial position is black or white, respectively, with the mean response to a gray bar in that position subtracted (Fig. 5C). In simple cells, presentation of white and black bars at the same spatial location is associated with opposite-sign modulation of firing rate relative to the firing rates elicited by gray bars, yielding values close to 1. In complex cells, either change in luminance elicits the same response, causing the firing rates to sum and leaving the PRFC near 0. The distribution of PRFC values (Fig. 5D) shows a high prevalence of complex cells and reflects a bimodal distribution often seen in previous V1 cell classifications (Skottun et al., 1991; Mechler and Ringach, 2002; Priebe et al., 2004). Comparing the model-based PRM and model-free PRFC, we find significant agreement between the two measures (Fig. 5E; also compare 5B and 5C), further validating our model-based results.

As a direct application of these measures, we calculate the degree of nonlinearity across cortical layers, expecting increasing degrees of nonlinearity outside the input layers of V1 (layer IV). The PRFC measure of phase-invariance displays the expected distribution across cortical layers, with cells in layer IV having much lower values on this indicator of phase-invariance (Fig. 6A) (Yeh et al., 2009; Sharpee et al., 2011; Atencio et al., 2012). As expected, simple cells (Fig 6C, D; left column) being already well described by a linear filter, are not much improved by an increasingly nonlinear model, while complex cells are poorly described by the LN model and thus show great improvement by the PCNIM relative to the LN model (Fig. 6C, D; right column). Additionally, we find that infra-granular cells display longer onset latencies on average compared to Layer IV cells (Fig 6B), as previously described (Nowak et al., 1995).

**Figure 6:**
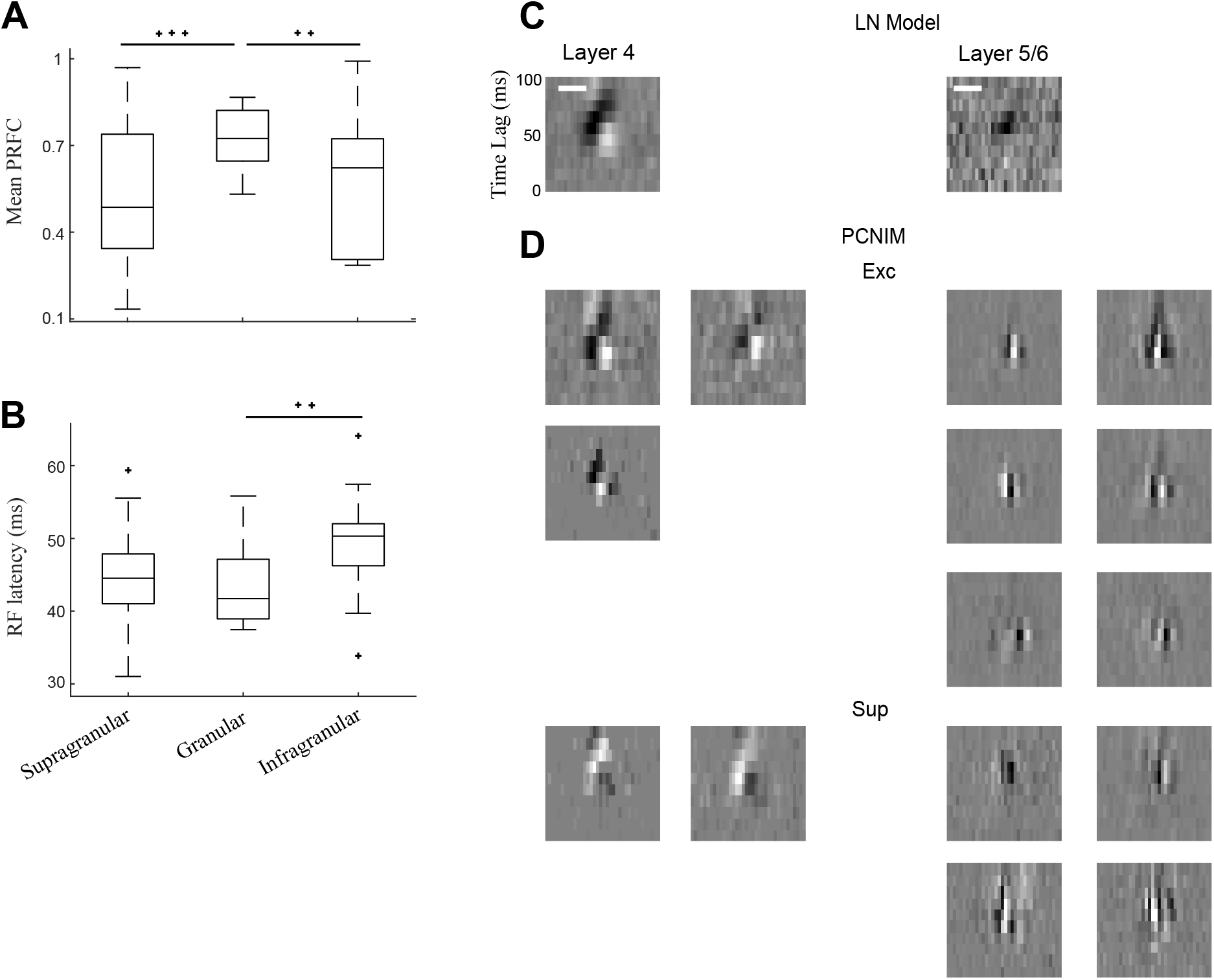
Degree of Nonlinearity across cortical layer. ***A***, PRFC results across the recorded population. Cells in Layer 4 have high PRFC values, while infra-and supra granular cells have low PRFCs, indicating a much higher proportion of complex cells in those layers, in agreement with previous literature. ***B***, Mean latency of excitatory filters for each model. Cells from infragranular layers have slower onsets, indicating that our models capture a delay in processing onset. C ***and D***, Example LN (in **C**) and PCNIM (in **D**) fits for a layer 4 simple cell (left) and a layer 2/3 complex cell (right). Beyond having a greater number of filters and being able to fit the phase-invariant complex cell, the PCNIM reveals additional high-resolution RF structure in both cases.

## Discussion

We have demonstrated how model-based characterizations of V1 neurons offer a more sophisticated description that is consistent with – but goes beyond – classical measures of V1 selectivity. Indeed, the underlying motivation is that nonlinear statistical models that are fit to more complex spatiotemporal stimuli capture aspects of V1 selectivity that is not evident in their responses to simpler stimuli, and thus measures based on such models will necessarily provide more information. Model-based characterization bridges the gap between recent advances in statistical modeling of neuron responses to complex stimuli (Butts, 2019) and the descriptions of V1 tuning established by classical stimuli. Thus, they can relate what can be relatively uninterpretable, complex models, to the fundamental stimulus selectivity of the modeled neurons. As such, we this approach as a new benchmark for describing V1 neuron selectivity that is applicable to its processing of complex stimuli.

We have illustrated such measures, and their implications, using an established nonlinear modeling framework (McFarland et al., 2013). Much of our analysis relies on our model’s ability to identify as given neuron’s selectivity as a function of multiple subunits, as opposed to a single receptive field filter. Based on our analyses, most V1 neurons exhibit a diversity of subunits, in that individual cells are well described by many subunits that are often selective to multiple distinct spatiotemporal elements, and for example might have direction-selective subunits at the same time as those tuned to non-moving features. This not only further highlights the diversity of selectivity exhibited by V1 neurons, but is also an example of the strengths of our approach in detecting this variability.

Furthermore, the approaches presented here can be applied to any model of V1 whose first stage consists of multiple spatiotemporal processing elements and otherwise is image-computable (e.g., generates a predicted response to stimulus input), and thus in principle can be applied to the range of models from spike-triggered neural characterization (e.g., (Touryan et al., 2002, 2005; Rust et al., 2005)) to recent models based on deep neural networks (e.g., (Kindel et al., 2017; Klindt et al., 2017; Moskovitz et al., 2018; Cadena et al., 2019)). However, we expect that such characterizations will be most accurate in models that accurately capture the neural computations (i.e., have high predictive power) and spatiotemporal processing attributable to individual V1 neurons, and thus use the NIM (McFarland et al., 2013; Almasi et al., 2020) in this paper for the basis of these characterizations.

A key finding of our analyses is that multiple subunits conferred tuning to a broader range of stimulus properties than would have been observable under classical contexts using traditional analyses. In the case of spatial frequency tuning, we find that some cells, while capable of resolving high-resolution inputs as indicted by the SF tuning of their subunits, are also tuned to broader stimulation by low-resolution inputs that spread across the cell’s entire RF. This also potentially offers an explanation of how complex cells can still show some phase selective response patterns when presented with low-frequency gratings moving across the receptive field, while stationary gratings failed to elicit the same response (Movshon et al., 1978b). These findings are also consistent with more recent findings of envelope tuning (Zavitz and Baker, 2014; Gharat and Baker, 2017), in which V1 processing for boundary segmentation is described as a theoretical LNLN cascade-style model with high-frequency filters in the first LN step and a coarse-scale boundary determined by the second LN step. They similarly observed responses to high-SF stimuli when presenting contrast-reversing luminance gratings (Gharat and Baker, 2017), implying that the ternary bar stimuli used in this study likely drives V1 activity in similar ways. However, unlike their model, we did not observe a single filter outlining the envelope, but instead find that the response to the low-frequency envelope emerges out of the combined contribution of many more localized filters that are spatially scattered around the cell’s RF. Additionally, our model filters capture the high-resolution components of the cell RF as predicted by that study and thus provide a way of quantifying the differences between tuning to envelope and carrier frequencies of a visual signal. Because related theoretical work (Li et al., 2010) found that scattered high-frequency inputs may also be indicative of cells with broad orientation tuning, the interaction between subunit scatter and orientation tuning will need to be investigated in the future using more high-dimensional stimuli than those used in our study, such as bar stimuli presented at multiple orientations or 2-dimensional noise stimuli. Such an approach might also yield clearer insight into other tuning properties where we observed significant variability between subunits, such as direction selectivity.

Our model-based approach also allows us to investigate the degree of nonlinearity needed to capture the responses of different V1 populations despite the well-established diversity of nonlinearity and tuning properties of primary visual cortex (Mechler and Ringach, 2002; Goris et al., 2015; Talebi and Baker, 2016) at much higher spatial resolution than was previous possible. Our model-based measures demonstrate a bimodal distribution of the degree of nonlinearity (Chen et al., 2009; Mechler & Ringach, 2002), and allows for additional insight into the construction of phase-invariant RFs by showing the spatial extent of individual inputs and their overlap. Because of this increased spatial resolution, the utility of our models as a measure of phase-invariance thus also goes beyond what is possible with classical analyses, such as using relative modulation analysis after presenting grating stimuli (Kagan et al., 2002). A key point to emphasize is that it is only possible to study the effects of nonlinear interactions across space in the context of sufficiently complex stimuli. Our analyses of the resulting models could focus on interactions between filters at the same spatial position, which appear to be the real mark of whether the cell exhibits linear ‘simple-cell-like’ stimulus processing or spatial-phase invariant ‘complex-cell-like’ processing.

Likewise, more simple measures such as the number of subunits are not a reliable indicator of cell complexity (consistent with previous modeling studies (Rust et al., 2005; Park and Pillow, 2011; McFarland et al., 2013; Fournier et al., 2014; Cadena et al., 2019)). Indeed, we see that the smaller scale of spatial selectivity relative to the full RF of the neuron can require multiple filters regardless of whether neurons are phase-invariant. However, how this scale difference manifests depends on the assumptions of modeling: techniques such as spike-triggered covariance will often explain the “most variance” of the subunit ensemble with large quadrature pair filters (Rust et al., 2005; Touryan et al., 2005; Park et al., 2013) and leave finer elements of spatial selectivity to later filters – but such characterizations are consistent with small, localized filters (Lochmann et al., 2013). Indeed, statistical models that have flexibility in their nonlinear processing, which for example is afforded by rectified subunits (McFarland et al., 2013; Kriegeskorte, 2015; Butts, 2019) not only can better capture neural responses but offer a finer description of such nonlinear interactions.

We propose that our approach for model-based measures provides a new benchmark for quantitative estimation and comparison of tuning properties. The investigation of differences between neurons across cortical lamina is just one application of our approach – future experiments may make comparisons between other neural populations, such as across eccentricity or between C-O blobs, to probe the functional organization of the visual cortex by identifying which computational subunits change in different contexts. Furthermore, while we utilized relatively low-dimensional bar stimuli, the approach is applicable to responses to any complex stimuli, including more natural stimuli that vary in additional dimensions such as color and stimulus contrast. Indeed, while probing more stimulus dimensions requires more data to support the resulting models (Butts, 2019), models fit using those stimuli could reveal how such properties integrate with the selectivity measures described in this paper. Likewise, model-based characterizations might similarly be applied to “deeper” models with additional stages of nonlinear processing than the two LN stages we used here (Kriegeskorte, 2015; Moskovitz et al., 2018; Butts, 2019; Cadena et al., 2019), either to describe later regions in the visual system or to further explore the additional nonlinearities present in V1. Such deeper networks would also be a more powerful tool to accurately identify shared computation in the context of more complex, and thus more high-dimensional, stimuli. Finally, due to the structural similarities among sensory cortical areas, model-based characterization might be further adapted to auditory or somatosensory modalities, potentially providing insight into general processing patterns in cortex by allowing for more direct comparison of shared or distinct computational subunits.

---

**Figure A1:**
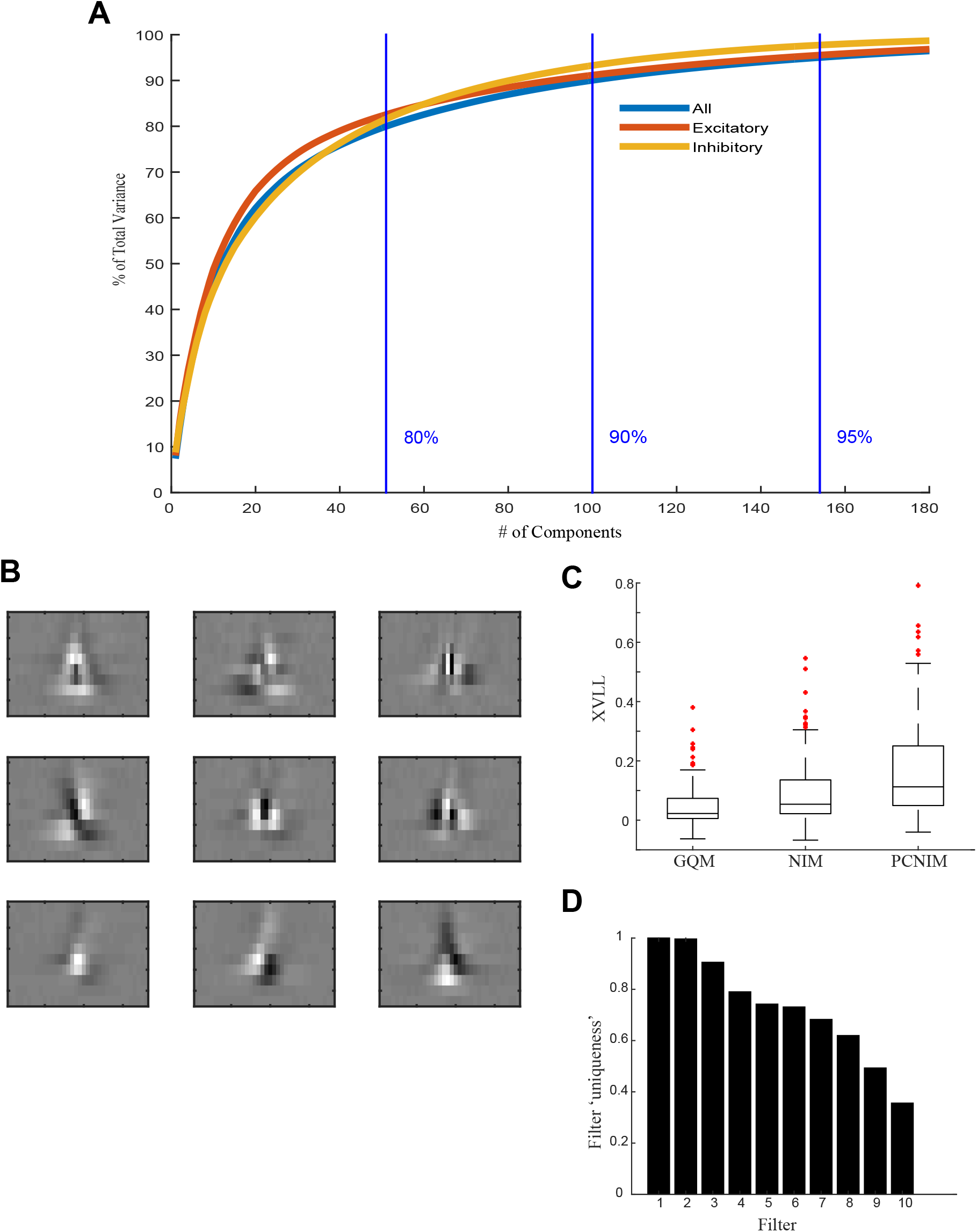
The Principal Component Nonlinear Input Model (PCNIM). ***A***, Variance explained by each Principal Component of a filter bank consisting of 1528 excitatory and 487 inhibitory filters. 80% of the total variance is explained by 51 PCs, 90% is explained by 100 PCs, and 95% is explained by 154 PCs. ***B***, First 9 Principal Components used for dimensionality reduction during modeling. ***C***, Log-Likelihood improvement of the models using the PC-space for dimensionality reduction relative to full-space models. The PCNIM clearly outperforms other models in describing the observed data. ***D***, ‘Uniqueness” of each filter of the example model shown in Figure 1D, quantified by the filter’s vector magnitude after orthogonalizing it from the previous filters. Most filters make significant contributions to outlining a different area of stimulus space the cell responds to.

